# Increasingly inbred and fragmented populations of *Plasmodium vivax* with declining transmission

**DOI:** 10.1101/100610

**Authors:** Andreea Waltmann, Cristian Koepfli, Natacha Tessier, Stephan Karl, Abebe Fola, Andrew W Darcy, Lyndes Wini, G. L. Abby Harrison, Céline Barnadas, Charlie Jennison, Harin Karunajeewa, Sarah Boyd, Maxine Whittaker, James Kazura, Melanie Bahlo, Ivo Mueller, Alyssa E. Barry

**Author notes:** Corresponding authors: Alyssa E. Barry and Ivo Mueller, Population Health and Immunity Division, Walter and Eliza Hall Institute of Medical Research, 1G Royal Parade, Parkville, Victoria, AUSTRALIA 3052. E OR.

## Abstract

The human malaria parasite *Plasmodium vivax* is resistant to malaria control strategies maintaining high genetic diversity even when transmission is low. To investigate whether declining *P. vivax* transmission leads to increasing *P. vivax* population structure that would facilitate elimination, we genotyped samples from a wide range of transmission intensities and spatial scales in the Southwest Pacific, including two time points at one site (Tetere, Solomon Islands) during intensified control. Analysis of 887 *P. vivax* microsatellite haplotypes from hyperendemic Papua New Guinea (PNG, n = 443), meso-hyperendemic Solomon Islands (n= 420), and hypoendemic Vanuatu (n=24) revealed increasing population structure and multilocus linkage disequilibrium and a modest decline in diversity as transmission decreases over space and time. In Solomon Islands, which has had sustained control efforts for 20 years, and Vanuatu, which has experienced sustained low transmission for many years, significant population structure was observed at different spatial scales. We conclude that control efforts will eventually impact *P. vivax* population structure and with sustained pressure, populations may eventually fragment into a limited number of clustered foci that could be targeted for elimination.

## Introduction

The international intensification of malaria control efforts over the last 15 years has reduced the global malaria burden by more than 50% with rapidly declining transmission in many endemic regions (WHO 2015a). *Plasmodium falciparum* and *Plasmodium vivax* are the major agents of human malaria, however, *P. vivax* is becoming the main source of malaria infection and disease in co-endemic areas because it is more resilient to control efforts (Harris *et al.* 2010; Kaneko 2010; Kaneko *et al.* 2014; Noviyanti *et al.* 2015; Oliveira-Ferreira *et al.* 2010; Rodriguez *et al.* 2011; Waltmann *et al.* 2015; WHO 2015b). These shifts in species dominance may result from the fact that *P. vivax* employs unique transmission strategies including: (i) dormant liver-stage infections (hypnozoites), (ii) pre-symptomatic development and continuous production of transmissible forms (gametocytes) during blood-stage infection, and (iii) lower density and thus less detectable infections. These biological characteristics suggest that *P. vivax* will be the far more challenging species to eliminate (Bousema & Drakeley 2011; Feachem *et al.* 2010; Mueller *et al.* 2013; White & Imwong 2012), and that interventions and monitoring approaches originally developed for *P. falciparum* malaria may not be sufficient or suitable for *P. vivax* (Alonso *et al.* 2011; Alonso & Tanner 2013; Cotter *et al.* 2013; Mendis *et al.* 2001).

Surveillance tools that monitor the impact of interventions are central to determining the success of disease control programs. Population genetics has been successfully harnessed to understand local changes in *P. falciparum* transmission dynamics in response to sustained control (Daniels *et al.* 2015), but this has not yet been applied extensively to *P. vivax*. Plasmodium parasites are haploid in the human host and replicate asexually for most of the lifecycle but undergo sexual replication and a brief period of diploidy within the mosquito vector. During this stage, meiosis produces haploid recombinant progeny (sporozoites) that are then inoculated back into the human host. The co-transmission of multiple clones to the vector is thus central to the generation and maintenance of genetic diversity via sexual recombination. As infections decline both within and among hosts, it is expected that effective population size, genetic diversity and gene flow will decrease, eventually leading to inbred, structured populations (Anderson *et al.* 2000a; Bousema *et al.* 2012; Markert *et al.* 2010). Conversely, in areas of high transmission, recombination between distinct clones and gene flow are more common, resulting in diverse, unstructured populations (Anderson *et al.* 2000a). Whilst *P. falciparum* fits this expectation (Anderson *et al.* 2000a), *P. vivax* populations retain high levels of diversity and large effective population sizes at low transmission (Barry *et al.* 2015; Chenet *et al.* 2012; Gray *et al.* 2013; Gunawardena *et al.* 2014; Van den Eede *et al.* 2010; Waltmann *et al.* 2015) and have higher diversity than *P. falciparum* populations (Jennison *et al.* 2015; Neafsey *et al.* 2012; Noviyanti *et al.* 2015; Orjuela-Sanchez *et al.* 2013). *P. vivax* population structure has been reported for some areas (Abdullah *et al.* 2013; Delgado-Ratto *et al.* 2016; Imwong *et al.* 2007; Van den Eede *et al.* 2010), but is absent in others (Jennison *et al.* 2015; Koepfli *et al.* 2013; Noviyanti *et al.* 2015). However, the population structure observed in these populations which include Peru (Van den Eede *et al.* 2010), Colombia (Imwong *et al.* 2007) and Malaysia (Abdullah *et al.* 2013), can be explained by multiple independent introductions of the parasite (Taylor *et al.* 2013), historically low *P. vivax* transmission (Delgado-Ratto *et al.* 2016; Van den Eede *et al.* 2010,), non-overlapping vector species refractory to non-autochthonous *P. vivax* strains (Pimenta *et al.* 2015) and historically focal transmission combined with recent reductions due to control (Abdullah *et al.* 2013). In regions with past hyperendemic *P. vivax* transmission and recent upscaling of malaria control efforts, population structure has not been observed (Noviyanti *et al.* 2015). The relationship between *P. vivax* transmission and population genetic parameters thus remains poorly understood, and requires systematic investigations with declining transmission and in the context of long-term intensified control.

Historically, the Southwest Pacific region, in particular PNG and Solomon Islands, has endured some of the highest *P. vivax* transmission anywhere in the world (Gething *et al.* 2012; Gething *et al.* 2011; Organisation 2013). This region has a natural, gradual decline in malaria endemicity from west to east with high transmission in PNG, moderate-to-high in Solomon Islands and low transmission in Vanuatu (Gething *et al.* 2012) that has been accentuated by recent control efforts (Hetzel *et al.* 2016; Koepfli *et al.* 2015a; Waltmann *et al.* 2015). Our previously published population genetic data from PNG and Solomon Islands (Jennison *et al.* 2015; Koepfli *et al.* 2013), combined with new samples from ongoing studies within Solomon Islands and Vanuatu, presents a unique opportunity to understand the population genetics of *P. vivax* in context with declining transmission. Here we have defined *P. vivax* population genetic structure at different transmission intensities, spatial scales and in the context of successful long-term malaria control. We analysed almost 900 *P. vivax* microsatellite haplotypes from isolates collected throughout the Southwest Pacific region (Figure 1), including dense spatial and temporal data from the Solomon Islands (Waltmann *et al.* 2015). The results demonstrate that *P. vivax* exhibits clear changes in population genetic parameters with declining transmission over space and time, highlighting the importance of maintaining control efforts, and the key role that population genetic surveillance of *P. vivax* can play in malaria control and elimination.

**Figure 1.**
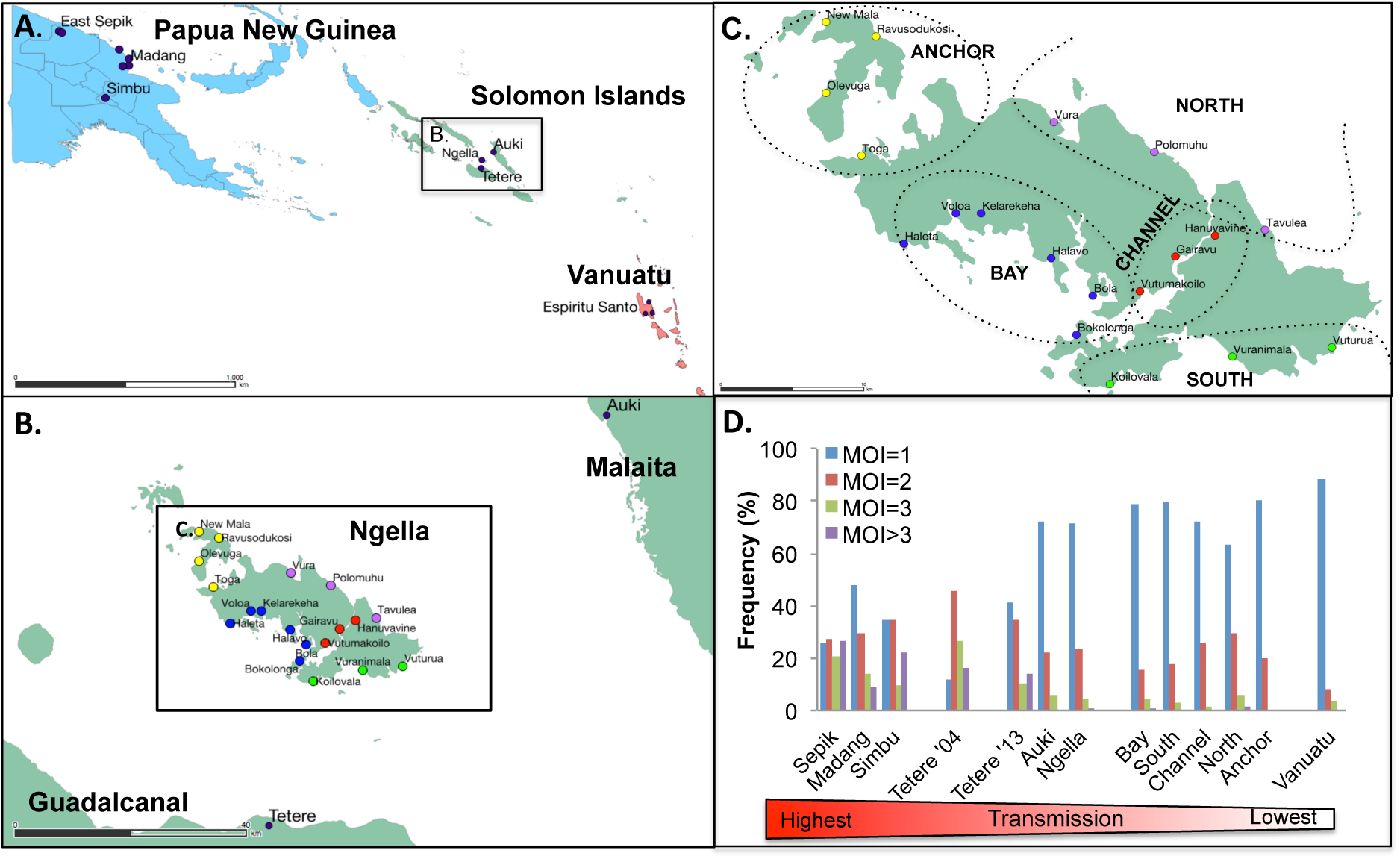
Map of the study areas and transmission intensity. (A) Southwest Pacific sampling locations showing Papua New Guinea in blue, Solomon Islands in green and Vanuatu in red, (B) central Solomon Islands, (C) Ngella, showing 19 villages and five distinct geographical/ecological regions. Anchor villages are indicated in yellow, Bay in blue, South Coast in green, Channel in red and North Coast in purple). (D) The distribution of multiplicity of infection values in each defined population, shown as an indicator of *P. vivax* transmission intensity.

## Results

### Wide range of *Plasmodium vivax* transmission intensities across the study area

Based on entomological inoculation rates and infection prevalence data, PNG, Solomon Islands and Vanuatu represent high, moderate to high, and low transmission areas respectively ((Gething *et al.* 2012), Figure 1, Table S1, Text S1). As reliable prevalence data was not available for all populations, we determined the multiplicity of infection (MOI) by genotyping of all available *P. vivax* infections using the highly polymorphic markers *MS16* and *msp1*F3, and calculated the proportion of polyclonal infections in each population as a surrogate measure of transmission intensity (Nkhoma *et al.* 2013). The distribution of complex polyclonal infections varied significantly across the Southwest Pacific (Figure 1D, Chi Squared test: p<0.0001) with polyclonal infections ranging from high in PNG (52.2%-74.3%), moderate to high in Solomon Islands (28.6-88.2%) to low in Vanuatu (12%)(Table S1). The Solomon Islands population of Tetere experienced a significant change in the distribution of polyclonal infections over a period of intensive control (Figure 1D, Chi Squared test: p = 0.0014) with polyclonal infections declining between 2004 (88.2%) to 2013 (58.6%, Table S1). There was significant variability also within both PNG and Solomon Islands (Chi Squared test: p<0.0001). In the Solomon Islands, Tetere 2013 (58.6%) had a higher proportion of polyclonal infections than Auki (28.6%) and Ngella (30.0%), consistent with lower transmission in the latter two regions. Within Ngella, an area of dense sampling divided into five distinct ecological zones (Anchor, North, Channel, South and Bay), the proportion of polyclonal infections ranged between 20.0-36.9% (Table S1), and was significantly associated with prevalence (Linear regression: r^2^ = 0.97, p=0.002).

### Definition of microsatellite haplotypes

Previously published microsatellite haplotype data was available for PNG (n=443) and the Tetere 2004 population (n=45, (Jennison *et al.* 2015; Koepfli *et al.* 2013)). New haplotypes were obtained for all monoclonal infections (MOI=1) from the Solomon Islands: Tetere 2013, Ngella and Auki populations, and from Vanuatu by genotyping an overlapping set of nine microsatellite markers. Two polyclonal infections (MOI=2) each from Auki and Vanuatu were also genotyped to boost sample numbers in those populations. Only high quality haplotypes with data for at least five out of nine well-validated microsatellite loci were retained for population genetic analysis (Jennison *et al.* 2015; Koepfli *et al.* 2013), resulting in seven samples being excluded. Two further haplotypes were identified as outliers (i.e. those that do not conform to the expected distribution), due to rare singleton alleles at the MS2 locus, and were discarded for subsequent analyses (data not shown). The final dataset comprised a total of 887 haplotypes including 443 from PNG, 420 from Solomon Islands and 24 from Vanuatu (Table 1, Dataset 1). Although most samples were initially identified as monoclonal, multiple alleles were detected after genotyping the additional nine markers. Thus the data includes 555 confirmed monoclonal infection haplotypes and 332 “dominant” haplotypes comprising the dominant allele calls (highest peaks) from samples with multiple alleles. The 887 haplotypes were distributed across all catchment areas, however smaller sample sizes were available for lower prevalence regions of Auki and Vanuatu (Table 1). Note that *MS16* and *msp1F3* were used only to determine MOI and are not recommended for analysis of population structure due to their extreme diversity (Koepfli *et al.* 2011; Sutton 2013).

**Table 1:** Genetic diversity of *Plasmodium vivax* populations of the Southwest Pacific.

| Country | Province | Population | *n* | *A*±SEM | *H*<sub>E</sub>±SEM | *R*<sub>S</sub>±SEM | *N*<sub>e</sub> (95% CI) | *N*<sub>e</sub> (95% CI) |
| --- | --- | --- | --- | --- | --- | --- | --- | --- |
|  |  |  |  |  |  |  | SMM | IAM |
| Papua New Guinea | East Sepik |  | 229 | 13.44±0.38 | 0.81±0.006 | 8.75±0.20 | 16059 (6901, 36582) | 6811 (2927, 15515) |
|  | Madang |  | 175 | 15±0.39 | 0.84±0.005 | 9.62±0.20 | 24293 (10439, 55337) | 8234 (3539, 18757) |
|  | Simbu |  | 39 | 8±0.46 | 0.81±0.011 | 7.37±0.38 | 15539 (6678, 35397) | 6713 (2885, 15291) |
| Solomon Islands | Guadalcanal | Tetere 2004 | 45 | 9.44±0.53 | 0.84±0.009 | 8.44±0.43 | 23220 (9978, 52893) | 8061 (3464, 18362) |
|  |  | Tetere 2013 | 39 | 7.78±0.3 | 0.79±0.009 | 7.05±0.24 | 11766 (5056, 26802) | 5963 (2562, 13583) |
|  | Central Islands (Ngella) | Bay | 82 | 12.33±0.27 | 0.81±0.006 | 9.20±0.15 | 16112 (6924, 36703) | 6821 (2931, 15538) |
|  |  | South | 35 | 9.11±0.34 | 0.82±0.012 | 8.50±0.32 | 16599 (7133, 37811) | 6911 (2970, 15744) |
|  |  | Channel | 47 | 9.33±0.35 | 0.79±0.009 | 8.10±0.28 | 11500 (4942, 26196) | 5907 (2538, 13456) |
|  |  | North | 136 | 13.89±0.28 | 0.85±0.004 | 9.73±0.17 | 26387 (11339, 60107) | 8564 (3680, 19509) |
|  |  | Anchor | 23 | 6.56±0.38 | 0.79±0.019 | 6.51±0.37 | 10771 (4629, 24535) | 5752 (2472, 13103) |
|  | Malaita | Auki | 13 | 5.33±0.54 | 0.80±0.026 | NA | 13170 (5659, 30000) | 6250 (2686, 14238) |
| Vanuatu | Sanma | Espiritu Santo | 24 | 5.56±0.34 | 0.72±0.022 | 5.45±0.33 | 4056 (1743, 9239) | 4131 (1775, 9411) |

*n*=number of microsatellite genotypes, *A* =number of alleles, *H<sub>E</sub>* = expected heterozygosity, *R<sub>s</sub>* = allelic richness based on smallest sample size, *N<sub>e</sub>* = effective population size, SEM = standard error of the mean, SMM = Stepwise mutation model, IAM = Infinite alleles model.

### Diversity and effective population size

Mean genetic diversity of the nine microsatellite markers showed a modest but significant trend of declining diversity from PNG (*H*_e_ = 0.81-0.84, *R*_s_ = 7.37-9.62) to Solomon Islands (*H*_e_ = 0.79 - 0.85, *R*_s_ = 6.51-9.20) and Vanuatu (*H*_e_=0.72, *R*_s_=5.45) (One way ANOVA test of trend: p <0.05, Table 1). There was a trend of decreasing diversity with declining polyclonal infections but this was not significant (*H*_e_, r^2^ = 0.23, p= 0.11, *R*_s_, r^2^ = 0.07, p = 0.42, Linear regression, Figure 2A,B). Effective population sizes (*N_e_*) reflect the high diversity across the different parasite populations. The Solomon Islands and PNG populations showed moderate to high *N_e_*, while Vanuatu had 1.5-5 fold lower *N*_e_ than any of the other populations (Table 1). The patterns observed suggest that low transmission, such as that seen in Vanuatu, is needed for significant reductions diversity and effective population size.

**Figure 2.**
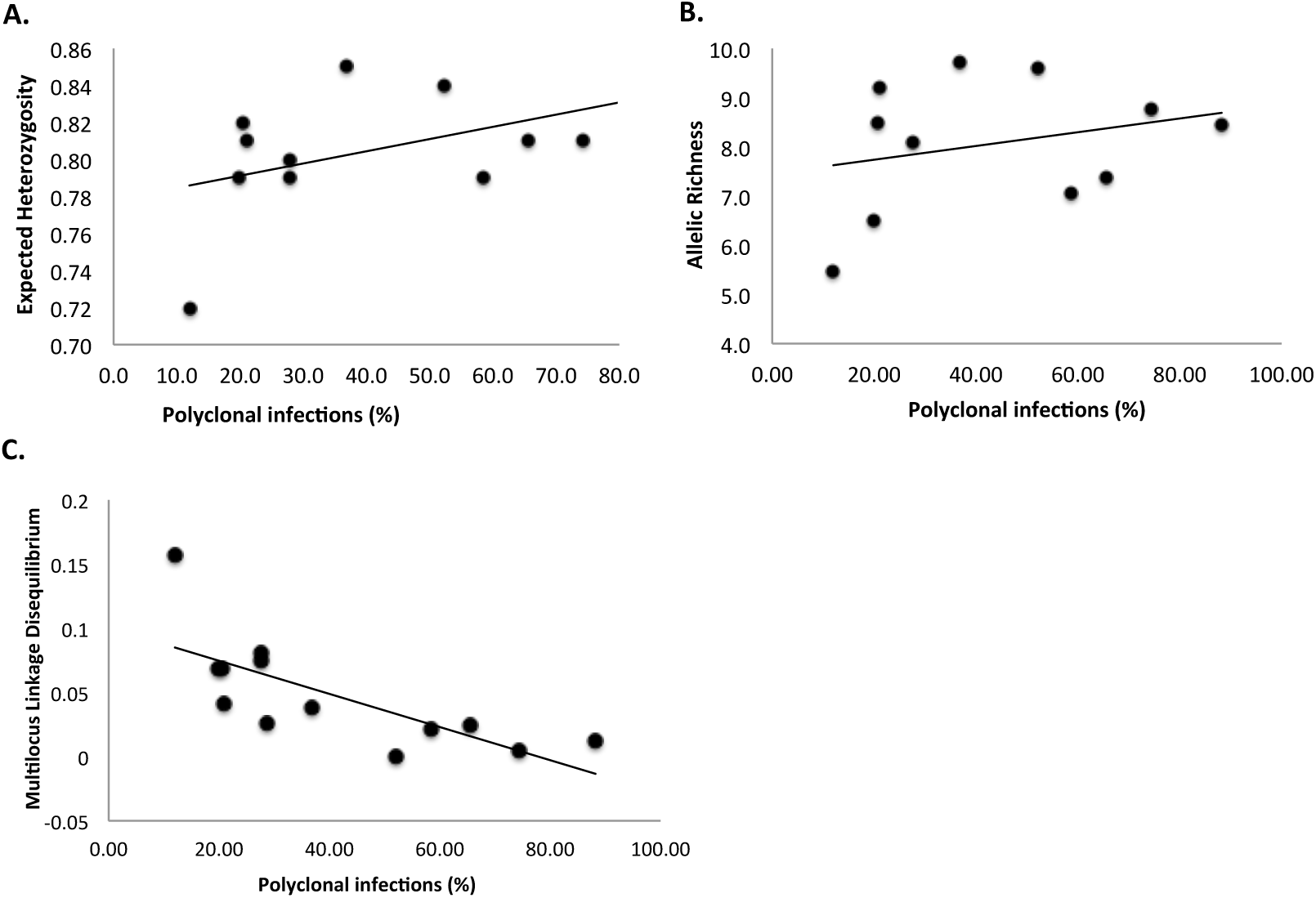
Relationship between transmission intensity and genetic diversity for *Plasmodium vivax* populations of the Southwest Pacific. Diversity of parasite populations based on the (A) mean expected heterozygosity (*H*_e_) and (B) Allelic richness (*R*_s_) and (C) multilocus linkage disequilibrium (*I*_A_^S^) was plotted against the proportion of polyclonal infections for each defined population (see Table 1 and S1 for values).

### Multilocus linkage disequilibrium

Multilocus linkage disequilibrium of asexual blood stage parasites provides an indirect measure of the rate of recombination between related individuals (inbreeding) in the mosquito stages, which is expected as transmission declines and infections become increasingly clustered. In previously published data from PNG and the Solomon Islands (Tetere 2004) there were no identical haplotypes and no significant LD was observed indicating limited inbreeding and random associations between alleles in those populations (Jennison *et al.* 2015; Koepfli *et al.* 2013). In the later data from Solomon Islands (i.e. Tetere 2013, Ngella and Auki) and Vanuatu, seven haplotypes were found repeatedly amongst 22 isolates, suggesting clonal transmission due to self-fertilization and no detectable recombination, or a single mosquito infecting several individuals. All repeated haplotypes were found in Ngella, and four were distributed among different villages or regions (Figure S1) making the latter scenario unlikely. Repeated and incomplete haplotypes were excluded for the analysis of multilocus LD retaining only the unique, complete microsatellite haplotypes comprised of all nine markers (n=248). Significant multilocus LD was observed in the Solomon Islands (Tetere 2013, Ngella and Auki) and Vanuatu populations (Table 2) (Koepfli *et al.* 2013). The pattern of multilocus LD was retained when only single infections were considered (n=93, Table 2), as well as when only one locus per chromosome was analyzed to confirm that LD was not the result of physical linkage (Table S2). Multilocus LD was significantly associated with the proportion of polyclonal infections (Linear regression: r^2^=0.55, p=0.005, Figure 2C).

**Table 2.**
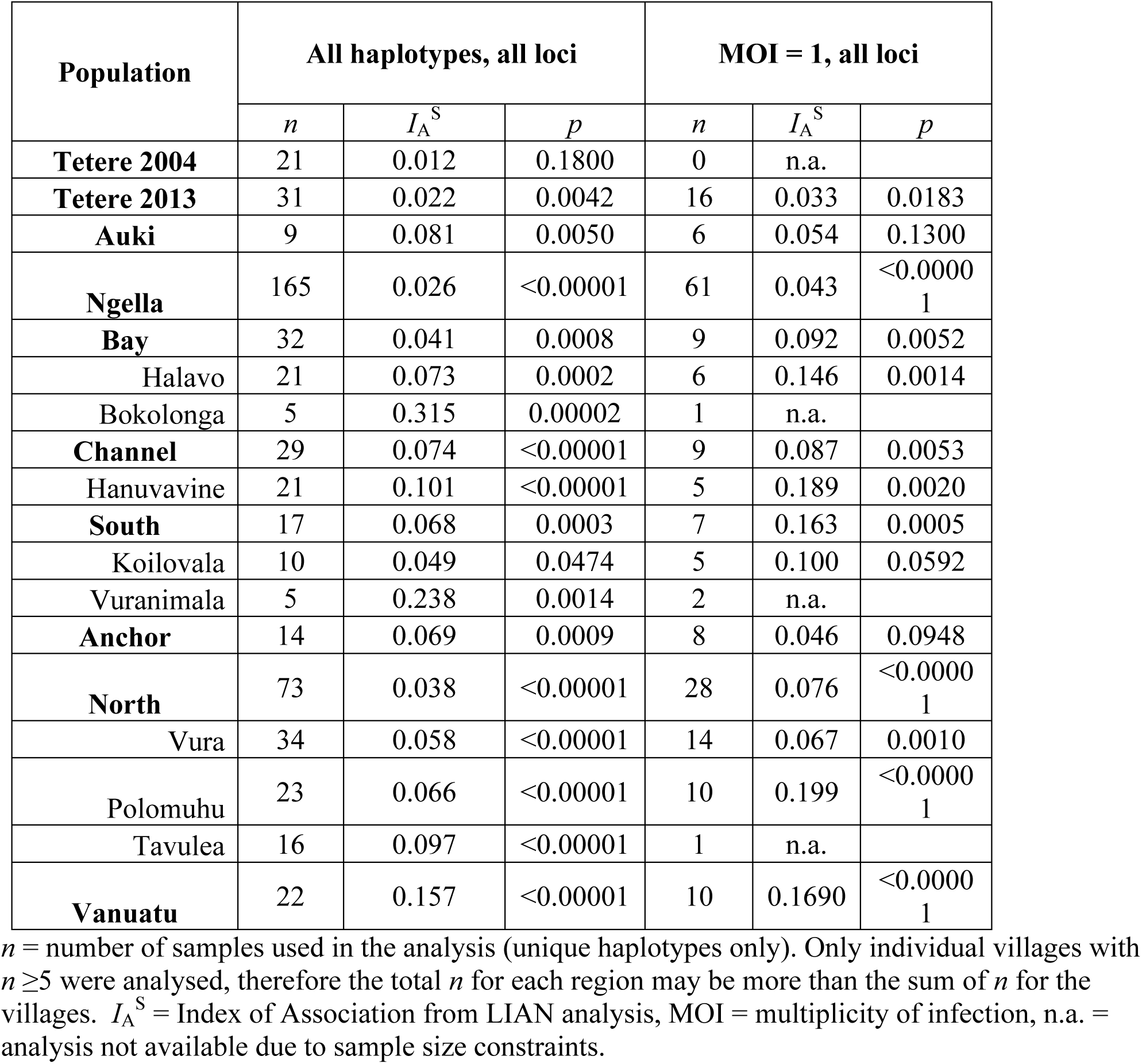
Estimates of Multilocus Linkage Disequilibrium (LD) in *Plasmodium vivax* populations of the Southwest Pacific. Multilocus LD values from PNG and Tetere 2004 populations are published elsewhere (Koepfli, et al. 2013; Jennison 2015).

| Population | All haplotypes, all loci |  |  | MOI = 1, all loci |  |  |
| --- | --- | --- | --- | --- | --- | --- |
| | $n$ | $I_A^S$ | $p$ | $n$ | $I_A^S$ | $p$ |
| **Tetere 2004** | 21 | 0.012 | 0.1800 | 0 | n.a. |  |
| **Tetere 2013** | 31 | 0.022 | 0.0042 | 16 | 0.033 | 0.0183 |
| **Auki** | 9 | 0.081 | 0.0050 | 6 | 0.054 | 0.1300 |
| **Ngella** | 165 | 0.026 | <0.00001 | 61 | 0.043 | <0.00001 |
| **Bay** | 32 | 0.041 | 0.0008 | 9 | 0.092 | 0.0052 |
| Halavo | 21 | 0.073 | 0.0002 | 6 | 0.146 | 0.0014 |
| Bokolonga | 5 | 0.315 | 0.00002 | 1 | n.a. |  |
| **Channel** | 29 | 0.074 | <0.00001 | 9 | 0.087 | 0.0053 |
| Hanuvavine | 21 | 0.101 | <0.00001 | 5 | 0.189 | 0.0020 |
| **South** | 17 | 0.068 | 0.0003 | 7 | 0.163 | 0.0005 |
| Koilovala | 10 | 0.049 | 0.0474 | 5 | 0.100 | 0.0592 |
| Vuranimala | 5 | 0.238 | 0.0014 | 2 | n.a. |  |
| **Anchor** | 14 | 0.069 | 0.0009 | 8 | 0.046 | 0.0948 |
| **North** | 73 | 0.038 | <0.00001 | 28 | 0.076 | <0.00001 |
| Vura | 34 | 0.058 | <0.00001 | 14 | 0.067 | 0.0010 |
| Polomuhu | 23 | 0.066 | <0.00001 | 10 | 0.199 | <0.00001 |
| Tavulea | 16 | 0.097 | <0.00001 | 1 | n.a. |  |
| **Vanuatu** | 22 | 0.157 | <0.00001 | 10 | 0.1690 | <0.00001 |

$n$ = number of samples used in the analysis (unique haplotypes only). Only individual villages with $n \geq 5$ were analysed, therefore the total $n$ for each region may be more than the sum of $n$ for the villages. $I_A^S$ = Index of Association from LIAN analysis, MOI = multiplicity of infection, n.a. = analysis not available due to sample size constraints.

### Population structure

To measure population structure across the study area, patterns of genetic differentiation among populations and local phylogenetic clustering of haplotypes was investigated. Average F-statistics over all loci indicated the presence of low levels of population subdivision amongst countries (*F*_ST_=0.049) and a gradient of increasing structure from high to low transmission. Negligible differentiation was observed among provinces in PNG (East Sepik, Madang and Simbu: *F*_ST_ =0.013), low levels among Solomon Islands provinces (Tetere 2013, Auki, Ngella: *F*_ST_ =0.035), and slightly higher genetic differentiation was observed among Ngella regions (Bay, South, Channel, North, Anchor: *F*_ST_=0.042). Very high genetic differentiation was found among the three Vanuatu villages (Port Orly, Nambauk, Luganville: *F*_ST_=0.348), however sample sizes were much smaller for this country (n per village=7-10, Figure 3A). Despite dense sampling within the Ngella regions, sample sizes were too small for village-level analysis within all but the North Coast region where it was similar to that found among the five Ngella zones (*F*_ST_=0.045)(Figure 3A). Pairwise Jost’s *D* statistics, which account for the high diversity of microsatellites (Jost 2008), confirm moderate to high differentiation among countries with 21.6-41.7% private alleles (Figure 3B). Within Solomon Islands, moderate to high proportions of private alleles were observed for Ngella:Channel (21.2-31.2%) and Auki (26.5-39.6%) compared to other populations. In addition, there was moderate genetic differentiation between villages on the Ngella:North Coast (18.4-23.6%) and high differentiation between Channel villages (48.8%)(Figure 3B). *G*_ST_ and *F*_ST_ values are also provided for comparison (Table S3).

**Figure 3.**
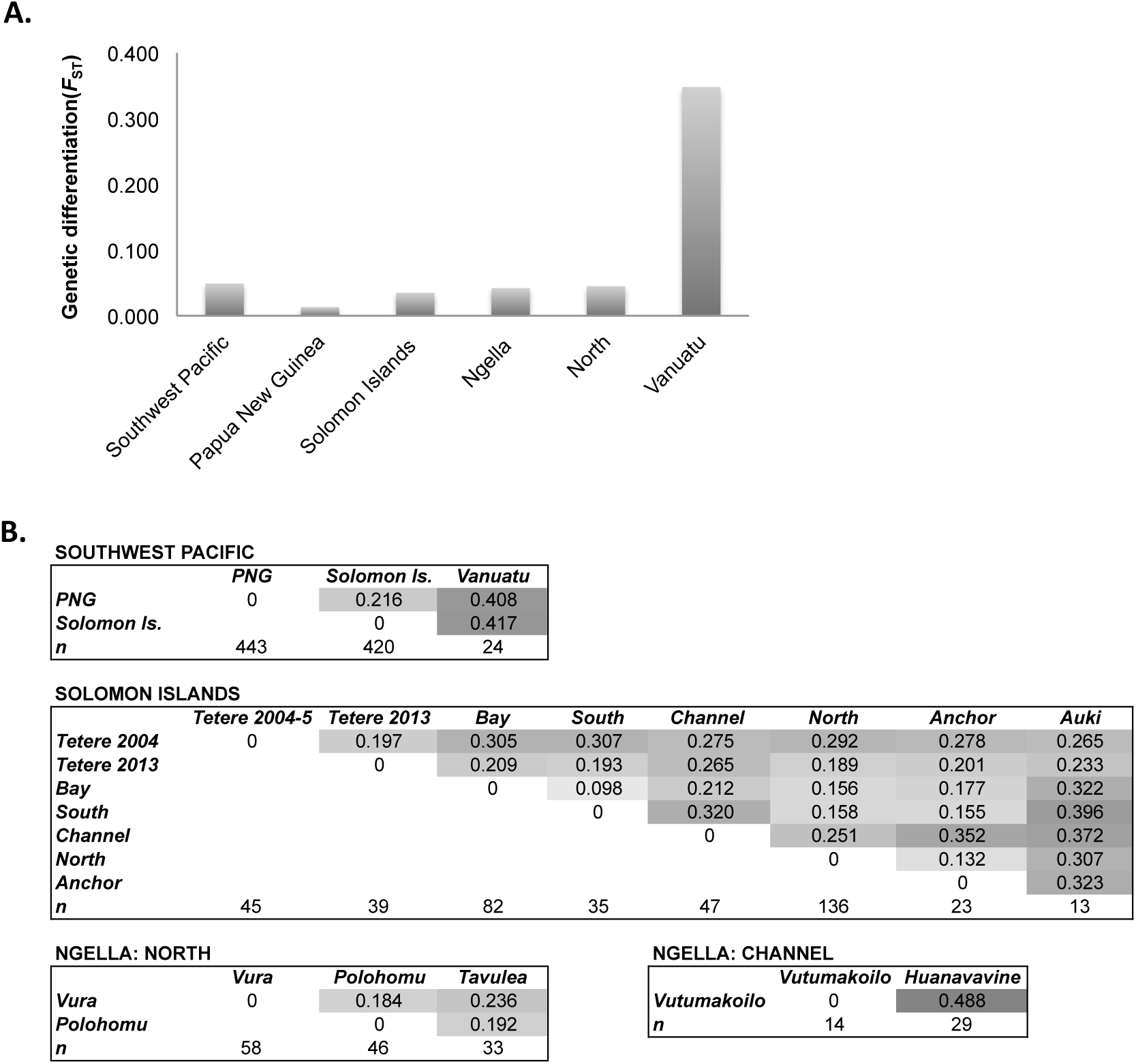
Genetic differentiation of *Plasmodium vivax* populations across the Southwest Pacific. (A) Average genetic differentiation among subpopulations. Average F statistics (*F*_ST_) was measured over all loci for all regions with at least two sub-populations of 20 or more samples, with the exception of Vanuatu, which had three populations of 7-10 samples. (B) Pairwise genetic differentiation between subpopulations. Pairwise differentiation was measured using Jost’s D, which accounts for the high diversity of microsatellite markers (Jost 2008). Values are shown for populations at different spatial scales. Darker shading indicates higher values.

To investigate population clustering patterns, we used the program STRUCTURE for K=1-20 to define genetic clusters within the entire dataset, as well as for Solomon Islands and its sub-regions. The analysis identified sub-populations at various spatial scales down to the village level (Figure 4, S2). Southwest Pacific parasites optimally cluster into three genetically distinct populations (K=3) associated with each of the countries (Figures 4, S3). For Solomon Islands, further substructure was observed at K=4, and within Ngella: Channel, with the Hanuvavine and Vutumakoilo village isolates forming distinct clusters; and on the North Coast, with some genetic clustering observed amongst villages (Figure 4, S2).

**Figure 4.**
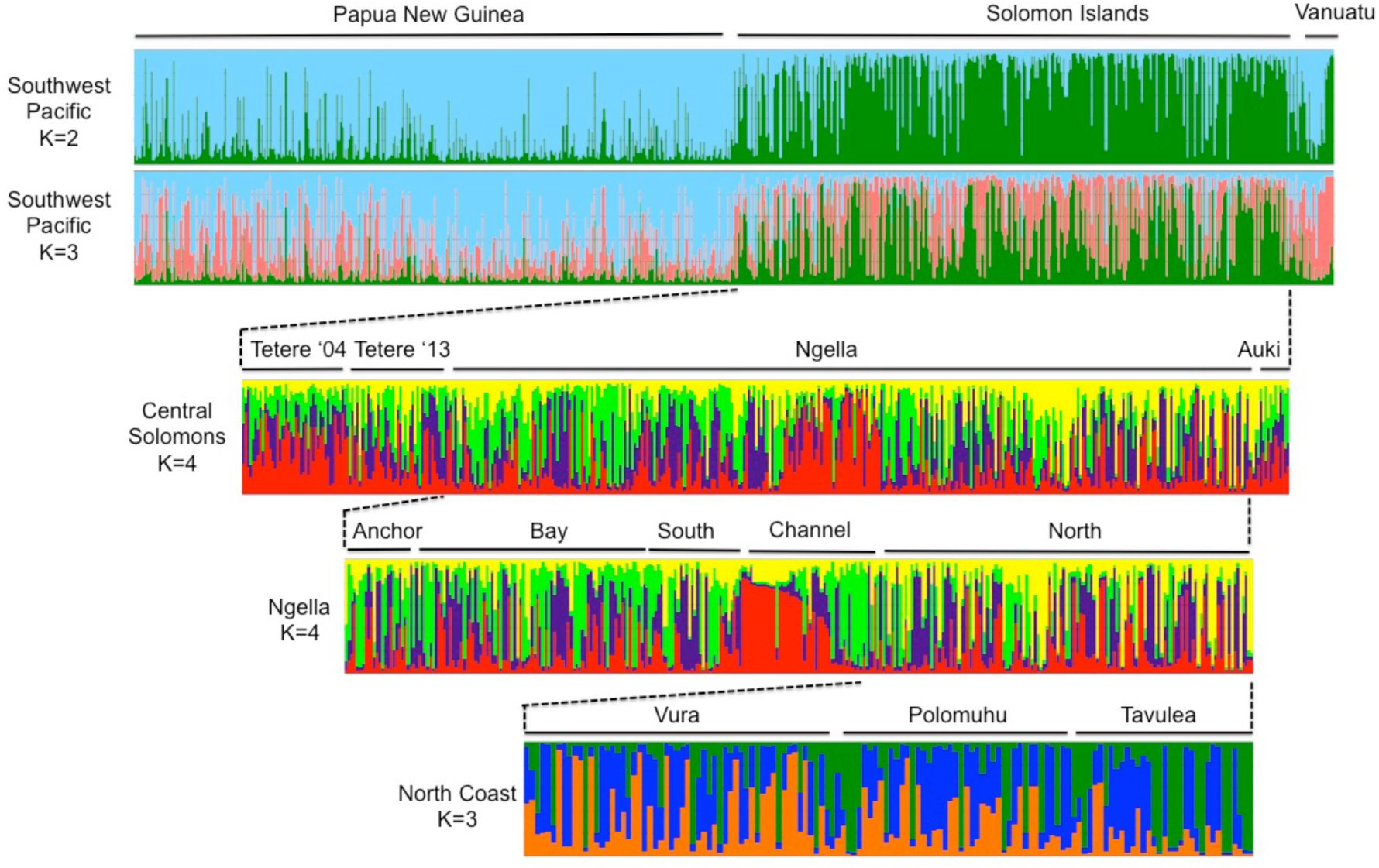
Geographical population clustering of *Plasmodium vivax* isolates of the Southwest Pacific. Results of STRUCTURE analysis are shown for different geographic strata including (A) Southwest Pacific, (B) Solomon Islands, (C) Ngella and (D) Ngella: North Coast. The analysis assigns *P. vivax* haplotypes to a defined number of genetic clusters (*K*) based on genetic distance. Vertical bars indicate individual *P. vivax* haplotype and colours represent the ancestry co-efficient (membership) within each cluster. Numbers indicate the haplotype ID.

Phylogenetic trees support the spatial structuring of haplotypes in Solomon Islands (Ngella) and in Vanuatu (Figure 5). High levels of recombination in *Plasmodium* lead to large star-shaped trees for large sample sizes, and thus only when isolates are closely related is clustering observed. In Ngella, the North Coast and Bay haplotypes radiate from distinct internal branches of the tree. Two distinct clusters were also observed for Channel isolates, one of which contains a number of closely related haplotypes and falls within a clade of North Coast isolates, the other with Bay isolates (Figure 5A). In Vanuatu, haplotypes clustered predominantly by village of origin (Figure 5B).

**Figure 5.**
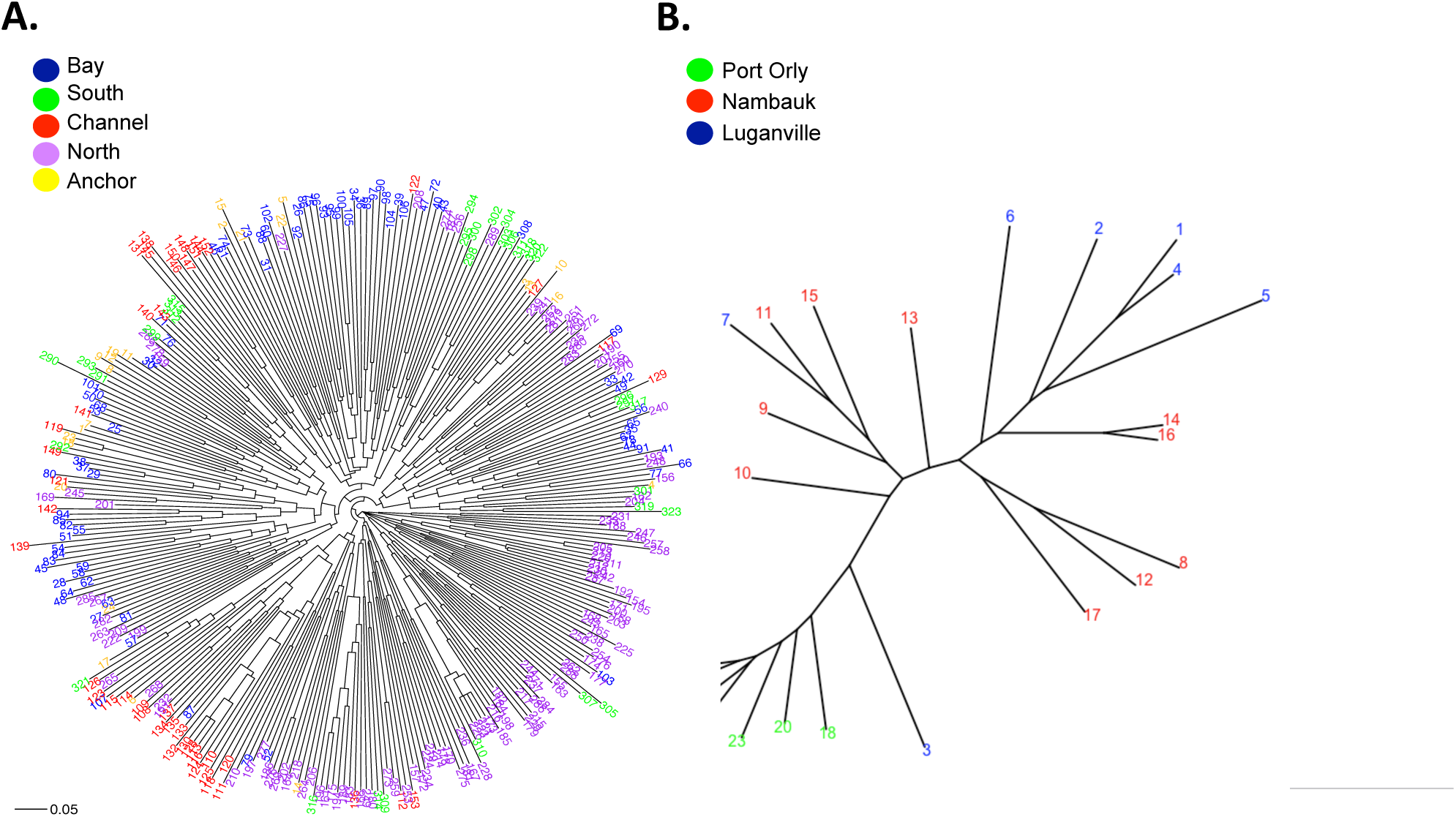
Phylogenetic analysis of *Plasmodium vivax* isolates of the Southwest Pacific. For the lower transmission regions of (A) Ngella and (B) Vanuatu, relatedness amongst haplotypes was defined by calculating the pairwise distance and visualized by drawing unrooted phylogenetic trees using the APE package in R software. Colours indicate the geographic origin of each sample as indicated in the key.

### Changes in *Plasmodium vivax* population structure during sustained control

The availability of samples from two time points during intensive malaria control in the Solomon Islands population (Tetere 2004 and 2013) allowed the investigation of changes in *P. vivax* population genetic parameters due to antimalarial intervention. In 2004, genetic diversity was high (*H*_e_=0.84, Table 1) and there was no significant multilocus LD (Koepfli *et al.* 2013) (Table 2). By 2013 however, diversity was significantly lower (*H*_e_ = 0.79, p=0.01, unpaired t-test), effective population sizes halved (Table 1), and multilocus LD increased to significance (Table 2). There were also low but significant levels of genetic differentiation between the two years (Jost’s D =19.7%, Figure 2B, *F*_ST_ = 0.029, Table S2). This suggests significant changes in the population structure of *P. vivax* in Tetere due to intensified malaria control.

## Discussion

As countries move towards elimination, measuring the impact of control efforts and changing transmission dynamics on parasite population structure will guide the switch from broad ranging to targeted control efforts, and will help prioritize specific geographic regions for elimination (Auburn & Barry 2017; Barry *et al.* 2015). Classical epidemiological studies that define the prevalence of infection are valuable as a monitoring tool, but only population genetic analyses such as those described here can detect the perturbation of transmission dynamics and population networks (Barry *et al.* 2015). Moreover, understanding the geographic distribution of malaria parasite populations including identification of inbred and fragmented populations is increasingly important as transmission drops to aid in the spatial stratification of interventions (Bousema *et al.* 2012). However, tracking the impact of control on *P. vivax* populations may be challenging given its more stable transmission at low prevalence, allowing populations to maintain high levels of diversity and gene flow (Hupalo *et al.* 2016; Jennison 2015; Noviyanti *et al.* 2015; Pearson *et al.* 2016). Using the largest and most densely sampled dataset of *P. vivax* genotypes to date, across a geographic region with a strong, natural gradient of transmission intensities, our results reveal that with declining transmission, there is a modest decrease in diversity but increasing multilocus LD and population structure. Observations in the Solomon Islands population of Tetere reveal a similar pattern with declining transmission due to malaria control. Together the results suggest that sustained control efforts are needed to reduce *P. vivax* diversity and interrupt gene flow. This provides a strong incentive to maintain intensive control efforts for *P. vivax* for longer periods relative to *P. falciparum*.

Despite reductions over time and space, relatively high genetic diversity and high effective population sizes remain in Solomon Islands *P. vivax* populations, however high multilocus LD and population structure was detected. High *P. vivax* genetic diversity at low transmission was first recognized in Sri Lanka (Gunawardena *et al.* 2014) and has also been observed together with significant multilocus LD in Peru (Chenet *et al.* 2012), Malaysia (Abdullah *et al.* 2013) and Indonesia (Noviyanti *et al.* 2015). Despite a substantial range of transmission intensities, the genetic diversity observed for PNG and Solomon Islands were similar while hypoendemic Vanuatu had much lower levels of diversity, indicating that *P. vivax* transmission must be sustained at low levels for long periods before population-wide genetic diversity is impacted. Multilocus LD and population structure may however signal changes in transmission intensity much earlier. The presence of identical haplotypes shared among Ngella parasites and significant multilocus LD is consistent with considerable levels of clonal transmission and inbreeding due to increasingly clustered transmission of clonal or highly related (including relapsing) infections. In most endemic regions, identical *P. vivax* haplotypes are rare and were seen only at very low transmission in Central Asia where the *P. vivax* population is nearly clonal or at low transmission in the Amazon (Koepfli *et al.* 2015b). With sustained low transmission, opportunities for recombination between diverse strains are reduced, and thus multilocus LD and consequent population structure may indicate interrupted *P. vivax* transmission.

The patterns of population structure observed likely reflect the contribution of relapse and polyclonal infections to outcrossing and gene flow (Fola *et al.* 2017; Jennison 2015). Relapse has been shown to account for up to 80% of *P. vivax* infections in the high transmission setting of PNG (Robinson *et al.* 2015) and would be expected to be even higher in a low transmission area. For some time after a reduction in transmission, the re-activation of parasites from a pool of genetically diverse hypnozoites from numerous past infections provides opportunities for the exchange and dissemination of diverse alleles, thus sustaining genetic diversity in the population. However, as the hypnozoite reservoir is depleted, focal clusters of infection may be composed of recent infections and subsequent relapses with highly related strains (Bright *et al.* 2014), and would explain high diversity in the context of significant LD. Thus relapse is likely to sustain residual transmission and maintain diverse meta-populations with high evolutionary potential. Other biological characteristics of *P. vivax* that are likely to sustain transmission and resilience to intervention include the rapid and continuous gametocyte production coupled with efficient transmissibility at lower gametocyte loads that drives high rates of human-to-vector transmission (Boyd & Kitchen 1937; Jeffery & Eyles 1955) and the rapid acquisition of clinical immunity early in life and low density of infection (Mueller *et al.* 2013) that would lead to a larger population reservoir of asymptomatic carriers (Barry *et al.* 2015; Harris *et al.* 2010; Waltmann *et al.* 2015). However, unlike relapse, these do not fully explain the patterns of population structure that we have observed in the context of declining transmission.

Across the Southwest Pacific, diversity amongst *P. vivax* populations was predominantly partitioned by country of origin and likely reflects both restricted gene flow and high LD in Solomon Islands and Vanuatu. Southwest Pacific *P. vivax* populations have also been shown to be genetically distinct from other worldwide populations (Hupalo *et al.* 2016; Koepfli *et al.* 2015b; Pearson *et al.* 2016). Only minimal population structure between PNG and Solomon Islands was previously reported for the small number of samples collected in Tetere between 2004-5 (Jennison *et al.* 2015; Koepfli *et al.* 2013). The differences with these earlier studies may be attributed to the much larger sample size from multiple Solomon Islands locations including Ngella and Auki, which have lower transmission than the previously analysed population of Tetere. The intervening intensification of antimalarial interventions in the region and consequent decrease in transmission may have also influenced the observed population structure. This is supported by the comparison of two time points for Tetere (2004 and 2013) that reveal a decline in polyclonal infections, corresponding with lower diversity and effective population size and an increase in multilocus LD.

The genetic structure of malaria parasite populations has previously been investigated with *P. vivax* populations over large spatial scales (e.g. between countries or distant locations within countries) (Arnott *et al.* 2013; Ferreira *et al.* 2007; Imwong *et al.* 2007; Jennison *et al.* 2015; Koepfli *et al.* 2015b; Koepfli *et al.* 2013). The high-resolution analyses of *P. vivax* population structure in the central zone of Solomon Islands, a region spanning around 100 km, revealed population structure among different island provinces. Ngella *P. vivax* populations were found to have moderate levels of genetic differentiation from populations of the other island provinces. Ngella is well connected to the higher endemicity areas of the Central Solomon Islands zone, as a direct and popular shipping route exists between Guadalcanal (Tetere) and Malaita (Auki) Provinces via Ngella. This suggests that despite a significant level of human movement among these three provinces, importation of *P. vivax* cases into Ngella may be sufficiently reduced to impact *P. vivax* gene flow. Whilst within country population structure has not been observed previously in the Southwest Pacific, it has been found in Malaysia (Abdullah *et al.* 2013), where prevalence of *P. vivax* has traditionally been described as focal, and has recently reduced; and in South America (e.g. Peru (Delgado-Ratto *et al.* 2016; Van den Eede *et al.* 2010), Venezuela (Chenet *et al.* 2012), Colombia (Imwong *et al.* 2007) and Brazil (Ferreira *et al.* 2007) where *P. vivax* has likely been introduced multiple times with adaptation to local vectors likely to have resulted in founder effects and influenced gene flow (Taylor *et al.* 2013). At reduced transmission in an African setting, *P. falciparum* populations were shown to become increasingly inbred with genetic relatedness rapidly increasing within the first year of intensified control, however this would be dependent on the structure and effective population size at baseline (Daniels *et al.* 2015). Previous population genetic data were only available for the Solomon Islands population of Tetere, which confirms similar changes for *P. vivax* under the pressure of control.

In the densely sampled study area of Ngella, Solomon Islands, *P. falciparum* has almost disappeared due to ongoing control interventions, but significant *P. vivax* residual, mostly sub-clinical transmission remains (Waltmann *et al.* 2015). In this region, *P. vivax* parasite populations showed levels of diversity that were associated with prevalence and polyclonal infections, and were spatially structured among different zones and even villages within the same region. Structured parasite populations within Ngella (20-50km) were subdivided into four genetic clusters distributed unevenly among Anchor/Bay/South (combined), North Coast and the two villages in the Channel region. The Channel villages lay in an extensive mangrove system on both sides of a channel however the area has comparable prevalence and proportions of polyclonal infections to other Ngella areas, showing that the population structure is likely to be influenced by the ecology and isolation of this region. Population structure was also observed among neighbouring villages of the North Coast. Thus, *P. vivax* population structure in Ngella seems to be organized as a hierarchical island model, consisting of a metapopulation of several sub-populations (Slatkin & Voelm 1991). Although no pre-2013 samples were available from Ngella, evidence from malaria surveys indicate a 90% reduction in cases from 1992 to 2013 (Solomon Islands National Vector Borne Diseases Control Program, unpublished data), consistent with the pattern of population structure observed being a direct result of malaria control. Thus, sustained intervention may have resulted in the relatively inbred and fragmented parasite populations observed that indicate a critical turning point may be within reach. While *P. vivax* may be overall more resistant to control efforts than *P. falciparum* (Bousema & Drakeley 2011; Feachem *et al.* 2010; Mueller *et al.* 2013; White & Imwong 2012), long-term sustained malaria control will place parasite populations under substantial stress and may lead to fragmentation of parasite subpopulations.

In summary, the results demonstrate changing *P. vivax* population structure with declining transmission over space and to a limited extent, time. While the data indicate that transmission is high in the PNG populations, inbreeding and population substructure was observed at all spatial scales within Solomon Islands and Vanuatu, consistent with increasing recombination of related clones within populations and hampered gene flow between populations. The fact that these patterns are observed after documented transmission decline and that temporal observations in one area suggest that long term malaria control has led to these patterns, however further investigations are needed to confirm this. These findings have significant public health implications showing that albeit more resistant to control efforts than *P. falciparum* (Alonso & Tanner 2013; WHO 2015a), *P. vivax* populations may eventually become increasingly inbred and fragmented if control pressure is maintained over an extended period. These results emphasize the need for interventions to eliminate *P. vivax* be sustained for very long periods, well beyond the time frame required for *P. falciparum* (Oliveira-Ferreira *et al.* 2010). Given the proposal to eliminate malaria from the Asia-Pacific by 2030 (APLMA 2014), intensive control pressure must be maintained to capitalize on these successes and avoid rebound. Enhanced control efforts including targeted control of transmission foci will help to reach these goals.

## Materials and Methods

### Study sites and *Plasmodium vivax* isolates

The *P. vivax* isolates used in this study are from PNG, Solomon Islands and Vanuatu. PNG has traditionally had the highest burden of the three countries and control has only been intensified in the last ten years (Hetzel *et al.* 2012). In Solomon Islands, sustained and intensified malaria interventions in the last two decades have resulted in an approximately 90% reduction in malaria incidence (Harris *et al.* 2010; PacMISC 2010; Waltmann *et al.* 2015; WHO 2015a). The small country of Vanuatu harbors the southern boundary of malaria transmission in the Pacific, as it is crossed by the Buxton Line, which defines the limit of Anopheline breeding (Maguire *et al.* 2006) resulting in a very low clinical infection rate that is dominated by *P. vivax* infections (WHO 2015a). At the time of sampling, transmission ranged from high in PNG (prevalence = 17.0-31.7%), moderate-high in Solomon Islands (3.9-31.7%) and low in Vanuatu (<1% (Gething *et al.* 2012), Table S1).

Genotyping data from total of 887 *P. vivax* isolates from PNG (n=443), Solomon Islands (n=420) and Vanuatu (n=24) were obtained (Table 1, Dataset 1). Samples were from both clinical and asymptomatic infections collected during different epidemiological surveys (Table S1). Data included previously published genotyping data from PNG collected in 2005-6 (n=443) and Solomon Islands in 2004 (Tetere 2004-5, n=45) (Jennison *et al.* 2015; Koepfli *et al.* 2013) in addition to 420 newly typed *P. vivax* isolates from multiple sites in the Solomon Islands collected in 2012-2013 (Waltmann *et al.* 2015)), and 24 genotypes from one site in Vanuatu collected in 2013 (Figure 1A). Dense sampling of the central region of the Solomon Islands allowed analyses at a wide range of spatial scales including three neighbouring island provinces including Guadalcanal (Tetere, n=39), Malaita (Auki, n=13) and Central Province (Ngella, n=323) (Figure 1B). In Ngella, sampling included 19 villages organized into five geographically and ecologically distinct areas including Bay (n=83), South (n=35), Channel (n=46), North (n=136) and Anchor (n=23, Figure 1C). In Vanuatu, samples were collected from the province of Espiritu Santo and included the villages of Port Orly (n=7), Luganville (n=7) and Nambauk (n=10). Further details of the samples and study sites are summarised in Table S1 and Text S1.

The study was approved by The Walter and Eliza Hall Institute Human Research Ethics Committee (12/01, 11/12 and 13/02), the Papua New Guinea Institute of Medical Research Institutional Review Board (11-05), the Papua New Guinea Medical Research Advisory Committee (11-06), the Solomon Islands National Health Research Ethics Committee (12/022) and the Vanuatu Ministry of Health (19-02-2013).

### Multiplicity of Infection (MOI)

To determine multiplicity of infection (MOI) in each population and to allow the selection of low complexity infections for the population genetics analyses, MS16 and *msp1*F3 genotyping data were used (Jennison *et al.* 2015; Koepfli *et al.* 2013). These data were previously available for the PNG (Jennison 2015; Koepfli *et al.* 2013), Tetere 2004 (Koepfli *et al.* 2013), and Ngella datasets (Waltmann *et al.* 2015). The MOI in the Tetere 2013, Auki 2013, and Vanuatu *P. vivax* populations was determined for this study, and was done according to previously published protocols (Jennison *et al.* 2015; Koepfli *et al.* 2013).

### Multilocus microsatellite genotyping

All confirmed monoclonal infections (MOI = 1) were then genotyped with nine genome-wide and putatively neutral microsatellites loci (MS1, MS2, MS5, MS6, MS7, MS9, MS10, MS12 and MS15) (Karunaweera *et al.* 2008). Due to small sample size, Auki and Vanuatu populations were supplemented by genotyping low complexity polyclonal infections (MOI=2). A semi-nested PCR strategy was employed, whereby a multiplex primary PCR was followed by nine individual secondary reactions, with a fluorescently labelled forward primer, as previously described (Jennison *et al.* 2015; Koepfli *et al.* 2013). PCR products were sent to a commercial facility for GeneScan^TM^ fragment analysis on an ABI3730xl capillary electrophoresis platform (Applied Biosystems) using the size standard LIZ500.

### Data analysis

Electropherograms resulting from the fragment analysis were visually inspected and the sizes of the fluorescently labeled PCR products were scored with Genemapper V4.0 software (Applied Biosystems), with the peak calling strategy done as previously described (Jennison *et al.* 2015). Raw data from the published dataset was added to the new dataset and binned together to obtain consistent allele calls. Automatic binning (i.e. rounding of fragment length to specific allele sizes) was performed with Tandem (Matschiner & Salzburger 2009). After binning, quality control for individual *P. vivax* haplotypes and microsatellite markers was conducted to confirm the markers were not in linkage disequilibrium (LD) and to identify outlier haplotypes and/or markers (i.e. haplotypes or markers which are disproportionately driving variance in the dataset). For isolates with an MOI=2, the dominant alleles were used to construct dominant clone haplotypes as previously described (Jennison *et al.* 2015).

Allele frequencies and input files for the various population genetics software programs were created using CONVERT version 1.31. Allele frequencies and genetic diversity parameters including the number of alleles (*A*), expected heterozygosity (*H_E_*) and allelic richness (*R_S_*) were calculated using *FSTAT* version 2.9.3.2. Effective Population Size (*N_e_*) was calculated using the stepwise mutation model (SMM) and infinite alleles model (IAM), as previously described (Anderson *et al.* 2000a). Mutation rates for *P. vivax* were not available and thus the *P. falciparum* mutation rate was used (Anderson *et al.* 2000b). For SMM, N_e_ was calculated as follows:

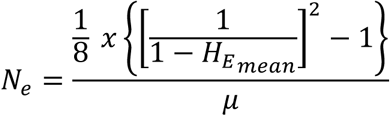

where *H*_E mean_ is the expected heterozygosity averaged across all loci.

For the IAM, *N*_e_ was calculated using the formula:

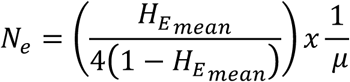

As a measure of inbreeding in the populations studied, multilocus LD (non-random associations between alleles of all pairs of markers) was estimated using the standardized index of association (*I*_A_^S^) in LIAN version 3.6. *I*_A_^S^ compares the observed variance in the number of shared alleles between parasites with that expected under equilibrium, when alleles at different loci are not in association (Haubold & Hudson 2000). The measure was followed by a formal test of the null hypothesis of LD and *p*-values were derived. Only unique haplotypes with complete genotypes were used and Monte Carlo tests with 100,000 re-samplings were applied (Haubold & Hudson 2000). The number of unique haplotypes was assessed using DROPOUT (McKelvey & Schwartz 2005). To confirm that LD was not artificially reduced by false reconstruction of dominant haplotypes, the analysis was also performed for the combined dataset of dominant and single haplotypes, and for single infections only. MS2 and MS5 both localize to chromosome 6 and MS12 and MS15 to chromosome 5 thus, analyses were repeated on datasets where MS5 and MS15 were excluded (chosen due to a greater degree of missing data) using the remaining seven loci spanning seven chromosomes. Where sample size permitted (n > 5), multilocus LD was also estimated at the village level.

To investigate geographic population structure, for each metapopulation we measured the weighted average F-statistics over all loci using the distance method (Weir & Cockerham 1984) using the global AMOVA implemented in Arlequin version 3.5.2.2 (Excoffier 2005). Pairwise comparisons among populations were done using three measures of genetic differentiation, namely *F*_ST_, *G*_ST_ and Jost’s *D*. *F*_ST_ was estimated using *FSTAT* version 2.9.3.2 (Goudet 1995). G_ST_ (Nei & Chesser 1983) and Jost’s *D* (Jost 2008) were estimated using the *R* package *DEMEtics*, as previously described (Gerlach *et al.* 2010). Population structure was further confirmed by Bayesian clustering of haplotypes implemented in the software STRUCTURE version 2.3.4 (Pritchard *et al.* 2000), which was used to investigate whether haplotypes cluster into distinct genetic populations (*K*) among the defined geographic areas. The analyses were run for *K*=1-20, with 20 independent stochastic simulations for each *K* and 100,000 MCMCs, after an initial burn-in period of 10,000 MCMCs using the admixture model and correlated allele frequencies. The results were processed using STRUCTURE Harvester (Earl & Vonholdt 2012), to calculate the optimal number of clusters as indicated by a peak in *ΔK* according to the method of Evanno *et al*. (Evanno *et al.* 2005). The programs CLUMPP version 1.1.2 (Jakobsson & Rosenberg 2007) and DISTRUCT 1.1 (Rosenberg 2004) were used to display the results. To assess phylogenetic clustering of haplotypes in each geographic area, the R software (APE) package was used to draw an unrooted phylogenetic tree using pairwise distances between multilocus haplotypes (Paradis *et al.* 2004).

Statistical analysis of epidemiological and population genetic parameters was done using Graphpad Prism version 7.

## Acknowledgements

The authors wish to acknowledge the support of the communities and field workers at all study sites. This study was supported by the International Centers of Excellence in Malaria Research (ICEMR, NIH grant U19AI089686) and by the National Health and Medical Research Council of Australia (NHMRC Project Grants 1021544, 1003825). IM is supported by a NHMRC Senior Research Fellowship (1043345) and AW was supported by an NHMRC Postgraduate Scholarship (#1056511). The authors acknowledge the Victorian State Government Operational Infrastructure Support and Australian Government National Health and Medical Research Council Independent Research Institute Infrastructure Support Scheme.

## Data Accessibility

The dataset is included in the manuscript submission (Supporting Files: Dataset 1), and will be submitted to DRYAD upon acceptance of the manuscript for publication.

## Author Contributions

AEB, JK and IM conceived the study, AW, CK, IM and AEB performed research, AW, AD, LW, HK, SB and MW conducted field studies, CK, GH, CB and CJ contributed data, AW, AF, SK, NT and MB analysed data, AEB, AW and IM wrote the paper, all other authors commented on and approved the manuscript.

## Supporting Information

**Figure S1.**
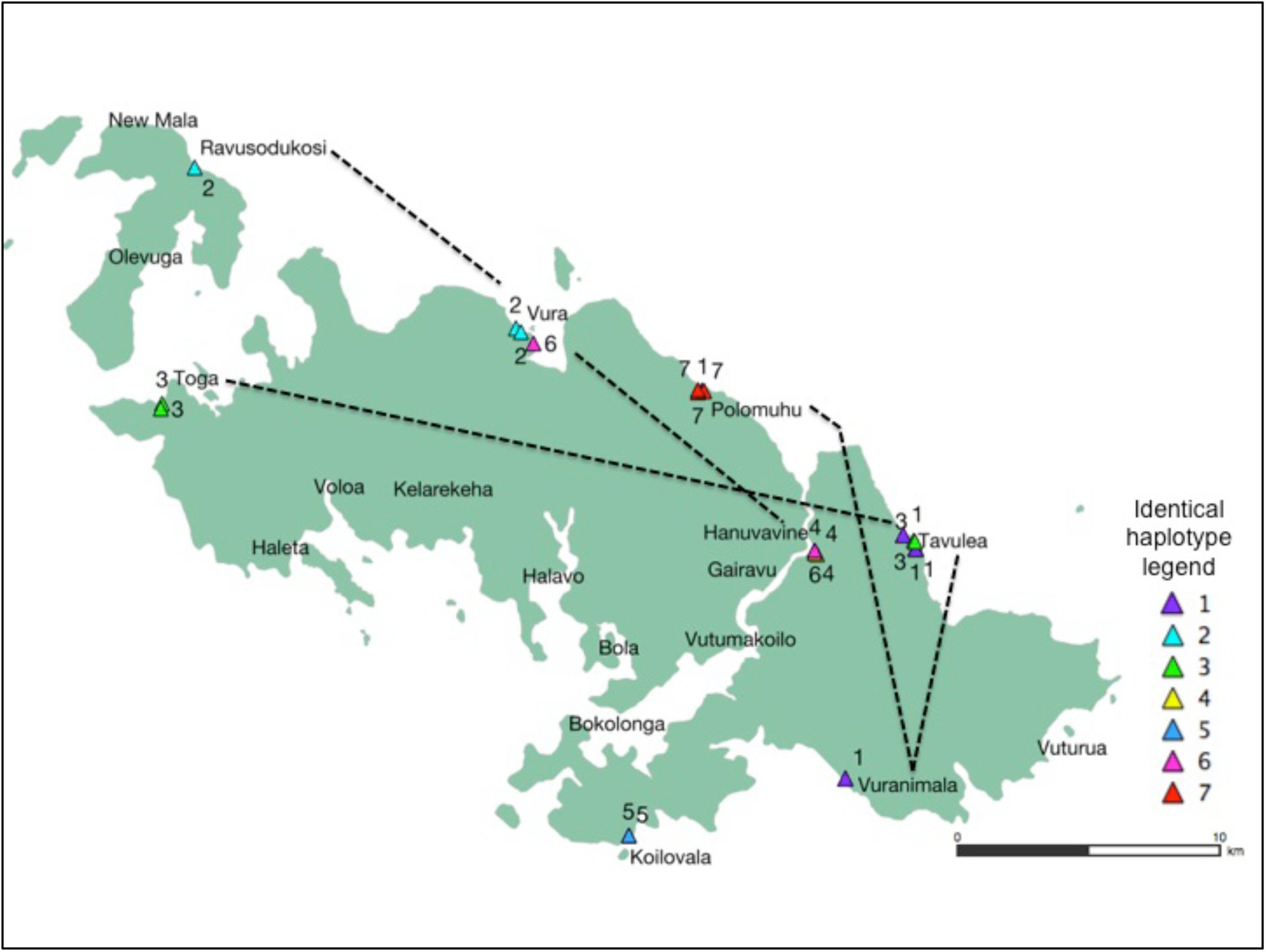
Spatial distribution of identical haplotypes across Ngella. Seven groups of identical haplotypes among 22 infections were identified. Identical haplotypes were found both within the same village (e.g. 4, 5, 7), and among villages and regions as denoted by dotted connectors (e.g. 1, 2, 3, 6).

**Figure S2.**
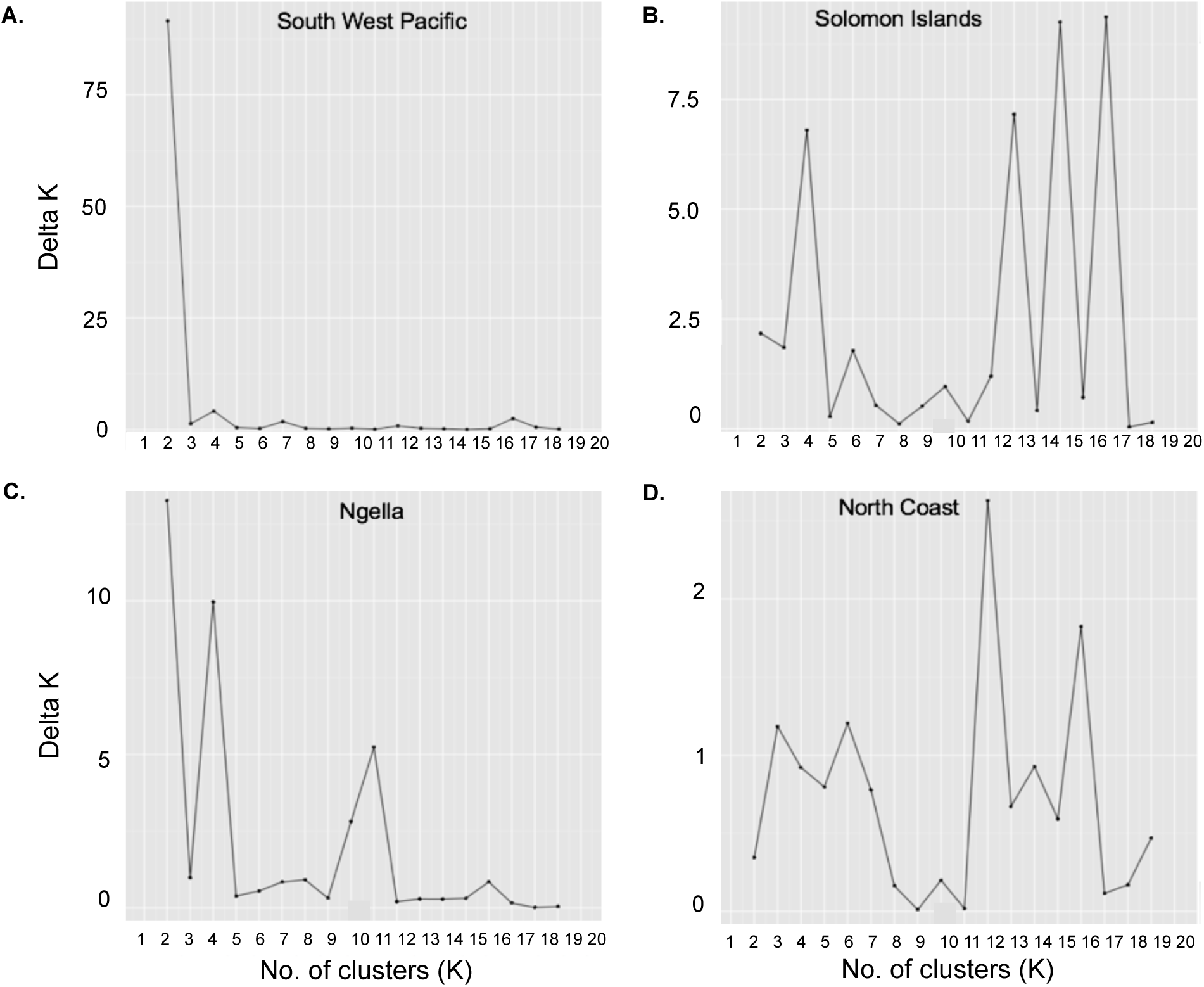
Definition of the optimum number of clusters for the STRUCTURE analyses. Peaks in Delta *K* (*ΔK*) identify the optimal of number of genetic clusters (*K*) and represent the uppermost hierarchical level of population structure. The first peak represents the optimal *K* identified and this value was used in interpreting the results of each of the respective analyses A-D. Subpopulation structuring may exist, which in our analyses is suggested by secondary peaks at higher K. (A) Southwest Pacific, K = 2, which was influenced by the small Vanuatu sample size (n=24) compared to the large sample size of PNG (n=443) and Solomon Islands (n=420). In this instance a K of 3, was determined by considering the uneven distribution of genetic clusters amongst countries. For (B) Solomon Islands and (C) Ngella, the optimal *K* was 4. For (D) North Coast villages of Ngella, the optimal *K* was 3.

**Table S1.** Study sites and sample details

| Country | Province | Catchment Area | Year | Survey type | *P. vivax* PCR prevalence (%) | No. samples genotyped for MS16/msp1F3<sup>c</sup> | Polyclonal infections (%) | Reference |
| --- | --- | --- | --- | --- | --- | --- | --- | --- |
| Papua New Guinea | East Sepik | Ilaita, Kunjingini, Wosera | 2003-2007 | Cross-sectional and longitudinal | 15-53<sup>a</sup> | 1350 | 74.3 | Lin et al. 2010, Koepfli et al. 2011 PLoS NTD, Arnott et al. 2013 |
|  | Madang | Alexishafen, Malala, Mugil, Utu | 2006 | Cross-sectional | 32<sup>b</sup> | 321 | 52.2 | Koepfli et al. 2013 PLoS ONE, jennison et al. PLoS NTD, Arnott et al. 2013 |
|  | Simbu | Sigimaru | 2004-2005 | Cross-sectional | n.a. | 48 | 65.7 | Koepfli et al. 2013 PLoS ONE |
| Solomon Islands | Guadalcanal | Tetere | 2004-2005 | Clinical | n.a. | 68 | 88.2 | Koepfli et al. 2013 |
|  |  | Tetere | 2013 | Clinical | n.a. | 56 | 58.6 | Wini *et al.*, unpublished |
|  | Central | Ngella Islands (all) | 2012 | Cross-sectional | 13.4 | 373 | 28.7 | Waltmann *et al.* 2015 |
|  |  | Bay |  | Cross-sectional | 13.7 | 105 | 21.0 | Waltmann *et al.* 2015 |
|  |  | South |  | Cross-sectional | 9.9 | 34 | 20.6 | Waltmann *et al.* 2015 |
|  |  | Channel |  | Cross-sectional | 13.2 | 54 | 27.8 | Waltmann *et al.* 2015 |
|  |  | North |  | Cross-sectional | 31.7 | 160 | 36.9 | Waltmann *et al.* 2015 |
|  |  | Anchor |  | Cross-sectional | 3.9 | 20 | 20.0 | Waltmann *et al.* 2015 |
|  | Malaita | Auki | 2013 | Clinical | n.a. | 18 | 27.8 | Wini *et al.*, unpublished |
| Vanuatu | Sanma | Espiritu Santo | 2013 | Clinical | n.a. | 25 | 12.0 | Karunajeewa *et al.*, unpublished |

<sup>a</sup>Data for Ilaita and Wosera only, <sup>b</sup>Data for Malala, Mugil and Utu only, <sup>c</sup>Number of samples with at least one marker successfully typed, n.a. = prevalence measurements not available as these samples were collected from confirmed *P. vivax* clinical cases.

**Table S2.** Estimates of multilocus linkage disequilibrium in *Plasmodium vivax* populations of the Southwest Pacific, considering only one locus per chromosome (excluding MS5 and MS15).

| Population | All samples, 1 locus/chromosome |  |  | MOI = 1, 1 locus/chromosome |  |  |
| --- | --- | --- | --- | --- | --- | --- |
| | n | $I_A^S$ | $p$ | n | $I_A^S$ | $p$ |
| **Tetere 2004-5** | 21 | 0.022 | 0.0976 | 0 | NA<sup>s</sup> |  |
| **Tetere 2013** | 31 | 0.018 | 0.0519 | 16 | 0.037 | 0.0435 |
| **Auki** | 9 | 0.082 | 0.0250 | 6 | 0.022 | 0.4230 |
| **Ngella** | 165 | 0.025 | <0.00001 | 61 | 0.040 | <0.00001 |
| **Bay** | 32 | 0.038 | 0.0081 | 9 | 0.087 | 0.0267 |
| Halavo | 21 | 0.071 | 0.0013 | 6 | 0.146 | 0.0084 |
| Bokolonga | 5 | 0.335 | 0.0002 | 1 | n.a. |  |
| **Channel** | 29 | 0.076 | 0.0001 | 9 | 0.087 | 0.0249 |
| Hanuvavine | 21 | 0.110 | 0.0001 | 5 | 0.176 | 0.0108 |
| **South** | 17 | 0.065 | 0.0036 | 7 | 0.130 | 0.0120 |
| Koilovala | 10 | 0.035 | 0.1330 | 5 | 0.057 | 0.3471 |
| Vuranimala | 5 | 0.264 | 0.0023 | 2 | n.a. |  |
| **Anchor** | 14 | 0.110 | 0.0002 | 8 | 0.019 | 0.3770 |
| **North** | 73 | 0.041 | <0.00001 | 28 | 0.090 | <0.00001 |
| Vura | 34 | 0.052 | 0.0000 | 14 | 0.091 | 0.0007 |
| Polomuhu | 23 | 0.070 | 0.0001 | 10 | 0.224 | <0.00001 |
| Tavulea | 16 | 0.107 | 0.0001 | 1 | n.a. |  |
| **Vanuatu** | 22 | 0.154 | <0.00001 | 10 | 0.189 | 0.00002 |

n = number of samples, $I_A^S$ = Index of Association from LIAN analysis, MOI = multiplicity of infection, n.a. = analysis not available due to sample size constraints.

**Table S3.** Pairwise estimates of genetic differentiation *G*_ST_ on the lower left; and *F*_ST_ on upper right)

|  | PNG | Solomon Islands | Vanuatu |
| --- | --- | --- | --- |
| PNG |  | 0.20 | 0.053 |
| Solomon Islands | 0.038 |  | 0.054 |
| Vanuatu | 0.085 | 0.085 |  |

**Solomon Islands**
|  | Tetere 2004 | Tetere 2013 | Ngella | Auki |
| --- | --- | --- | --- | --- |
| Tetere 2004 |  | 0.029 | 0.046 | 0.046 |
| Tetere 2013 | 0.021 |  | 0.024 | 0.024 |
| Ngella | 0.028 | 0.015 |  | 0.040 |
| Auki | 0.039 | 0.029 | 0.034 |  |

### Text S1. Additional information on study sites and samples

#### Papua New Guinea

Previously published microsatellite haplotype data were available from three areas Madang Province (n=175), East Sepik Province (n=229) and Simbu Province (Sigimaru, n=39) (Arnott *et al.* 2013; Jennison *et al.* 2015; Koepfli *et al.* 2013), and represent both asymptomatic and clinical samples collected during cross-sectional and cohort studies in 2005-2006 prior to the implementation of widespread intensified control. Full details of these samples are available in the relevant publications.

#### Solomon Islands

Malaria transmission in the Solomon Islands is highly heterogeneous throughout the country, with the highest number of cases in Central Islands, Guadalcanal and Malaita provinces (PacMISC 2010), the three of the most populous provinces (approximately 40% of national population (Government 2009)). In 2013, *P. vivax* accounted for 47.6% of all confirmed clinical cases of malaria nationwide and it was the predominant species circulating among asymptomatic individuals in the community (Harris *et al.* 2010; National Vector Borne Diseases Control Program 2013; Waltmann *et al.* 2015).

Solomon Islands *P. vivax* haplotypes originated from isolates collected in three provinces, namely Central Province (Ngella), Guadalcanal (Tetere) and Malaita (Auki). The three provinces are separated by sea (Figure 1), but are well connected by a frequent and popular ferry service and numerous, unscheduled motorized boat trips. Ngella is a group of three islands, namely Anchor, Big Ngella and small Ngella. Tetere is located 30 km east of the national capital Honiara, on Guadalcanal Island and Auki is the capital of Malaita province.

The Ngella *P. vivax* isolates were collected as part of a large household-based, cross–sectional survey of 3501 participants of all ages conducted in May-June 2012 (Waltmann *et al.* 2015). This survey demonstrated Ngella to be an area of considerable residual *P. vivax* (prevalence by PCR =13.4%, 468/3501), but disappearing *P. falciparum* transmission (prevalence by PCR =0.14%, 5/3501) (Waltmann *et al.* 2015).

Tetere isolates were collected at two time points during a period of intensified malaria control. Tetere 2004 isolates were collected from a clinical trial conducted between 2004-5 (Ballif *et al.* 2010). This microsatellite data was described elsewhere previously (n=45, (Koepfli *et al.* 2013)). During that time, Tetere was considered as high transmission, and the study which collected the isolates, reported a *P. vivax* prevalence of 19.1% by light microscopy (LM) (Ballif *et al.* 2010), a diagnostic method which is substantially less sensitive than the PCR methods used in the Ngella survey (Waltmann *et al.* 2015). Tetere 2013 isolates were clinical samples collected in the context of an antimalarial drug efficacy trial conducted in 2013 (Wini et al., unpublished).

Samples from Auki were also from a clinical trial conducted in 2013 (Wini *et al*., unpublished). Prevalence data was not available for the Tetere 2013 nor the Auki study.

#### Vanuatu

In 2013, a similar clinical trial was carried out on Espiritu Santo, with clinical samples being collected from participants residing in three villages Port Orly, Nambauk and Luganville (Boyd et al., unpublished). Prevalence data was not available for this area.

